# Can Bayesian phylogeography reconstruct migrations and expansions in human history?

**DOI:** 10.1101/2020.07.20.212043

**Authors:** Nico Neureiter, Peter Ranacher, Rik van Gijn, Balthasar Bickel, Robert Weibel

## Abstract

Bayesian phylogeography aims to reconstruct migrations in evolutionary processes. This methodological framework has been used for the reconstruction of homelands and historical expansions of various language families, but its reliability for language diversification research has remained unclear. We contribute to this discussion with a simulation study where we distinguish two types of spatial processes: *migration* and *expansion*. By migration we denote long-distance movement of whole populations, leaving their previous habitat empty. Expansions are small-scale movements of speakers or inclusions of new speakers into the language community, cumulatively contributing to a gradual spread into new territories. We simulate migrations, in the form of directional random walks, and expansions, in the form of a grid-based region-growing process. We run both simulation scenarios with varying degrees of directional trends and evaluate the performance of state-of-the-art phylogeographic methods. Our results show that phylogeography fails to reconstruct migrations, but works surprisingly well on expansions, even under severe directional trends. We demonstrate that migrations and expansions have typical phylogenetic and spatial patterns, which in the one case inhibit and in the other facilitate phylogeographic reconstruction. Furthermore, we propose descriptive statistics to identify whether a real sample of languages (Bantu), their relationship and spatial distribution, better fits a migration or an expansion scenario. Bringing together the results of the simulation study and theoretical arguments, we make recommendations for judging the adequacy of phylogeographic models to reconstruct the spatial evolution of languages.

## 1 Introduction

Is it possible to reconstruct the place of origin of a language family just by looking at the current geographic distribution of the extant languages? This certainly seems to be a difficult task. A common way to reconstruct the spread of cultural traits is through diffusion models [1, 2], originally applied in ecology [3], which use historical data to map out spatial dispersal over time. But in the absence of historical data, as is usually the case for the analysis of language spread, diffusion models would naively place the origin at the center of mass of the present-day locations.

As an alternative, it is sometimes proposed that the homeland of a language family lies in the area of maximal diversity, by analogy with patterns in biological evolution [4]. However, the mechanisms of language diversification are subject to additional social and environmental factors that challenge this model [5,6,7,8,9]. Also, there is no mechanistic link between rates of diversification and homeland location. We cannot exclude a slow-down of diversification rates caused by spatial saturation and stable co-existence in the homeland. Nor can we exclude a loss of diversity in the homeland due to later spreads. This makes inferences of homeland locations from diversity patterns unreliable.

Given these shortcomings of previous approaches, Bayesian phylogeography promises a clear improvement for reconstructing evolutionary processes: It incorporates the phylogenetic relationship between languages to inform the reconstruction of the spatial expansion. Indeed, Bayesian phylogeography has been used to reconstruct the spread of several language families, for example, Arawak [10], Indo-European [11], Bantu [12], Pama-Nyungan [13] or Sino-Tibetan [14]. Some of these reconstructions were corroborated by archaeological evidence and previous results [12, 13], while others challenged existing hypotheses [11]. This raises an obvious question: Do these models have the scientific authority to challenge established views on history?

Like all model-based inference, phylogeographic methods are based on assumptions. It is crucial to understand in what cases these assumptions are violated and whether such violations lead to errors in the reconstruction. We see a lack of literature discussing the ways in which Bayesian phylogeography works and what kind of spatio-temporal processes can actually be reconstructed. These issues are particularly important in the case of linguistic phylogeography, where historical data is usually scarce and ambiguous.

In this paper, we evaluate the adequacy of Bayesian phylogeographic methods for reconstructing the spatial evolution of languages. We simulate virtual processes based on different historical scenarios and then try to reconstruct the simulated processes using phylogeographic methods. Our evaluation metrics are based on the error between the reconstructed and simulated root location (i.e. homeland). Of course, the remainder of the process, what happened between the root and the tips, is of interest as well, but focusing on the root simplifies the quantification of the reconstruction error and is indicative of the quality of the whole reconstruction. Furthermore, reconstructing the homeland of a language family takes a very prominent role in many studies in historical linguistics (e.g. [10, 11, 15]).

A simulation-based evaluation has two main advantages over an empirical approach: 1) We can perfectly evaluate the reconstructions, since the exact simulated migrations are known. Such knowledge of a ground truth is scarce in real historical scenarios. 2) We can control all parameters of the process, and this allows us to draw conclusions about the conditions under which the reconstruction works.

We distinguish between two historical processes - migration and expansion - and for each of them we evaluate the reconstruction quality under varying degrees of directional trends. For the purposes of the simulation study, we define migration as a random walk process and expansion as a grid-based region-growing process. The importance of migration and expansion processes in human history has been a source of debate in archaeology and historical linguistics and is sometimes associated with entire migrationist and diffusionist research traditions that emphasize one process over the other [16, 17]. In this study, we examine the specific impact of these two processes on phylogeographical reconstruction success.

Historically, we understand **migration** as the permanent movement of entire populations to inhabit a new territory, separate from the one in which they were previously based. Potential causes for large-scale migrations include: Environmental push and pull factors, such as climatic changes, migrations of game or the search for more fertile lands. For example, **(author?)** [18] discuss Chadic migrations in the context of climatic changes at the end of the African humid period. Migration, as we describe it here, is a demic process, but there are cultural equivalents, where a language is pushed out of its homeland by another one through a process of language shift (and thus spreads to areas elsewhere). An example is the displacement of the Celtic languages from most of mainland Europe (except Brittany) to the British Isles. What is critical from the point of phylogeographical modeling is that the languages have completely left their homeland.

The term **expansion** describes a series of small-scale movements of people (also called demic diffusion or dispersal) or new people adopting the language (cultural diffusion), accumulating to a slow but steady spread of a language. A common explanation for major historical expansions is population pressure, where growing populations require more resources and territory. Specifically, this explanation has led to the proposal of the *farming/language dispersal hypothesis* [19], positing that “the present-day distributions of many of the world’s […] language families can be traced back to the early developments and dispersals of farming”. In this context, the spreading of many large language families has been described as expansions, including the Bantu, Arawak or Sino-Tibetan languages [19].

We further distinguish between language spread with and without a **directional trend**. While directional trends in migrations typically arise from an intrinsic bias in one direction, expansions also often show directional trends emerging from geographical or ecological constraints, such as oceans, mountains or deserts. If languages expand, but restrictions only allow an expansion in one direction, the resulting spread will follow a directional trend. The reconstruction of migrations or expansions under directional trends seems particularly challenging: How can we know where a directed movement of multiple languages over a long time period started if we only know their current locations? If the whole language family is involved in this movement, the reconstruction seems outright impossible. If only some of the languages were involved, the remaining (sedentary) languages might still uncover the true location of the homeland.

We have seen that both expansion and migration can be driven by demic or cultural processes. However, Bayesian phylogeographic modeling is blind to these processes: reconstruction is based entirely on the spatial patterns and the relationship between the languages. Thus, the distinction between demic and cultural processes is crucial in the discussion of the results, but it does not affect the design of the simulation study.

We implement the concepts of migration and expansion in two corresponding simulation scenarios, both with varying degrees of directional trends. We implement the migration simulation in the form of directional random walks. In a random walk process, particles (in this case representing languages) move in space stochastically. In the case of directional random walks, this stochastic movement has an intrinsic bias in one direction. This model has been used in movement analysis to study animal migrations [20, 21] or the spread of viruses [22].

We then further implement a simulation of historical linguistic expansions, with and without directional trend. In this scenario languages are represented by geographic areas, rather than point locations. These areas expand and once they reach a certain size the language splits up. This seems like a reasonable assumption, since large geographic areas make contact between some of the people in the population less likely and at some point, this leads to diversification [23]. After the split, the two resulting languages continue to grow and split separately. Geographical constraints and clashes with other populations force this expansion into one predominant direction.

We will show how these two approaches responsible for implementing the simulation of migration and expansion processes, respectively, lead to quite distinct spatial patterns, resulting in strong differences in the accuracy of the phylogeographic reconstruction.

## 2 Phylogeographic modeling

Relationships between languages have been modeled as trees since the 19th century, but the field is increasingly adopting phylogenetic methods from evolutionary biology to model language trees [24, 25, 26]. These methods reveal both the pattern of how organisms split, i.e. the topology of the phylogenetic tree, and the rate at which organisms split, i.e. the branch lengths in the tree. In addition, they allow the inference of trait evolution – the change of properties along the tree [27].

State-of-the-art methods of phylogenetic analysis rely on computational algorithms and statistical models, in particular Bayesian inference and Markov chain Monte Carlo sampling (MCMC) [28]. Bayesian phylogenetic analysis proposes possible evolutionary histories, and evaluates these against the likelihood of the data and prior beliefs. The analysis yields a posterior distribution and, thus, captures the inherent uncertainty of the evolutionary process [28]. There are several computer programs for Bayesian phylogenetic analysis, the most frequently used being BEAST [29] and MrBayes [30].

Bayesian phylogeography [31, 32] aims to infer the spatial expansion of a phylogenetic tree, that is, the spatial location of all ancestral nodes (including the root) and the rate at which organisms or languages change their location – the diffusion rate. The spatial locations make up the path of the expansion. The diffusion rate scales the expansion to the geographic and temporal context. Originally, Bayesian phylogeographic analysis was developed to reconstruct the spatial diffusion of viruses [31]. Recently, it has also been applied extensively in linguistics to explore the historic expansion of language families [11, 12, 10, 13, 14].

Phylogeography can be viewed as a special case of trait evolution, where the changing property is the spatial location, modeled as a continuous position on a plane [31, 10] or sphere [33]. We briefly outline how these models allow inferences and summarize their main properties.

Bayesian phylogeographic models accommodate a Brownian diffusion (BD) [34] process along the phylogenetic tree. Starting at a potential spatial location of the root, the BD moves down the branches of the tree, proposing locations for all ancestral nodes in turn, until reaching the observed tips [22]. We show samples from such a BD process along a fixed tree, ending in fixed tip locations in Figure 1 a) and the resulting distributions at the internal nodes in b). We can see that the uncertainty is 0 at the tip locations and increases as we move further back in time. In two-dimensional space, the likelihood of moving down a branch is modelled as a bivariate normal distribution with a displacement mean and variance. The mean is usually set to zero, i.e. there is no directional trend in the expansion. The variance is proportional to the rate of diffusion times the branch length, which is the amount of time needed to reach the next ancestral node. In an MCMC, possible spatial expansions are proposed and evaluated against their likelihood. Intuitively, a spatial expansion yields a high likelihood if it minimizes the distance from the root through all ancestral nodes to the observed tips. In addition to that, fossils – known spatial locations of extinct organisms (or languages, in our case) [35] – can help to place ancestral nodes, and thus, improve reconstruction.

**Figure 1:**
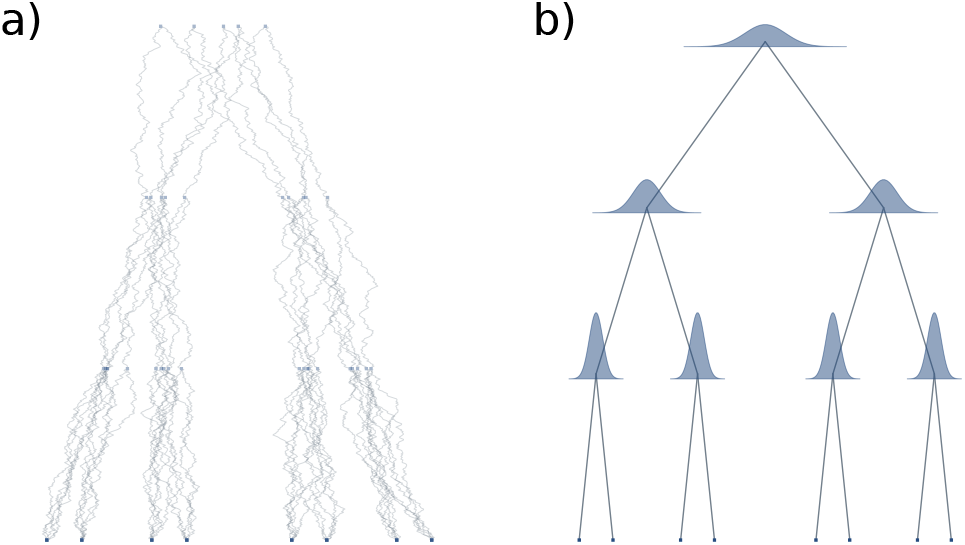
For a given phylogenetic tree different random walks can explain the data. a) Five examples of random walks (in one spatial dimension), leading to the same tip locations. b) Marginal posterior distributions of each internal node location under this random walk model.

There are three state-of-the-art phylogeographic models, all of which build on the concept of Brownian diffusion as outlined above:

- Random walk (RW) models assume the diffusion rate to be constant along the entire phylogeny. This implies that the expansion in space is homogeneous [31].
- Relaxed random walk (RRW) models allow the diffusion rate to vary and the dynamics of the expansion to change along the tree. In RRW, bursts of rapid migrations can be followed by phases where a population remains stationary [31]. Since a dynamic expansion is realistic in many application examples [12, 36, 31], RRW models are the de-facto standard for phylogeographic reconstruction.
- Directional random walk models allow the displacement mean to differ from zero. This introduces a directional trend (or directional bias), such that the expansion is free to prefer a particular spatial direction. The constant directional random walk (CDRW) [37] assumes the displacement mean to be constant over the entire expansion, whereas in the relaxed directional random walk (RDRW) it is allowed to vary [22]. In theory, directional random walk models should reconstruct directed migrations, but they rely on fossils to inform the inference [22].

In this article we evaluate the performance of phylogeographic models to correctly infer spatial patterns in language phylogenies resulting from either migrations or expansions, with particular attention to patterns with a directional trend. RW and RRW do not model a directional trend, suggesting that inference will favor a radial expansion from a location somewhere in the center of the observed distribution of the tips, even in the presence of a trend. While CDRW and RDRW models should be able to infer directional bias, it is unclear to what extent they allow the reconstruction of migrations or expansions without historical information in the form of fossils.

We evaluate the performance of phylogeographic reconstructions in a simulation study. In phylogenetics, simulation studies are widely used to validate the consistency of new models under model assumptions (e.g. [31, 22, 38]). In contrast to this, we simulate historically motivated scenarios of migrations and expansions, which intentionally violate some of the model assumptions. In that respect, our study is similar to [39], which explores the effect of horizontal transmission on phylogenetic reconstruction, a process not accounted for in basic phylogenetic inference.

## 3 Methods

We propose a simulation-based procedure to evaluate the performance of Bayesian phylogeographic methods, consisting of three steps:

1. *Simulate* a phylogenetic tree and movement of the languages in space.
2. *Reconstruct* the movement based on the simulated tree and tip locations using phylogeographic analysis.
3. *Evaluate* the results by comparing the reconstructed movements with the originally simulated ones.

In the first step we randomly generate phylogenetic trees and the spread of the corresponding languages in space and time. By changing the way in which we simulate the spread, we can test the sensitivity of the reconstruction to different movement scenarios. In particular, we are interested in the performance of phylogeographic methods in movement scenarios with directional trends. We explain the exact methodology behind each of these three steps in detail in the supplementary material (3-3). In this section we summarize the general set-up and terminology.

We simulate migrations (**MigSim**; supplementary material 3) in the form of directional random walks, where languages are represented by points moving stochastically in space. The directional trend is modelled as a constant bias in one direction. Languages split up into new languages at a constant rate, independent of the spatial movement. In contrast, expansions (**ExpSim**; supplementary material 3) are simulated in a region-growing process, where languages are represented by areas of cells on a grid, which expand to free neighboring cells at a constant rate. The areas split up and form new languages as they grow beyond a certain size. We limit the expansion to a circular sector of varying angles (representing geographical constraints), resulting in varying levels of directional trends. In the main model we do not allow language areas to overlap, but we relax this assumption in a sensitivity analysis (supplementary material 5.1). In order to compare the two simulation scenarios, we define a unified measure for the strength of the directional trend: The **observed trend** is the mean displacement from the root to the tip locations (supplementary material 3).

We use BEAST 1.10.4 [40] for a phylogeographic reconstruction of the homeland, based on the simulated phylogenies and tip locations (supplementary material 3). As evaluation scores we use the root mean square error (RMSE), bias (i.e. the systematic reconstruction error, ignoring errors due to variance in sampling and reconstruction) and highest posterior density (HPD) region coverage of the root location (supplementary material 3).

## 4 Results

We ran experiments based on the migration (MigSim) and expansion (ExpSim) scenarios. In both scenarios we look at cases with different levels of directional trends and we reconstruct the root using both the standard relaxed random walk (RRW) and the constant directional random walk (CDRW) models. The results for the MigSim and the ExpSim scenarios are presented in Figures 2 and 3, respectively. Figures 2a and 3a show the bias and root mean square error (RMSE) of the reconstruction, indicating how far we expect the reconstructed homeland to be located from the real one. Figures 2b and 3b depict the 80% and 95% HPD coverage, indicating the overconfidence (coverage is below 80%/95%) or underconfidence (coverage is above 80%/95%) of the model. In what follows we summarize the main results.

**Figure 2:**
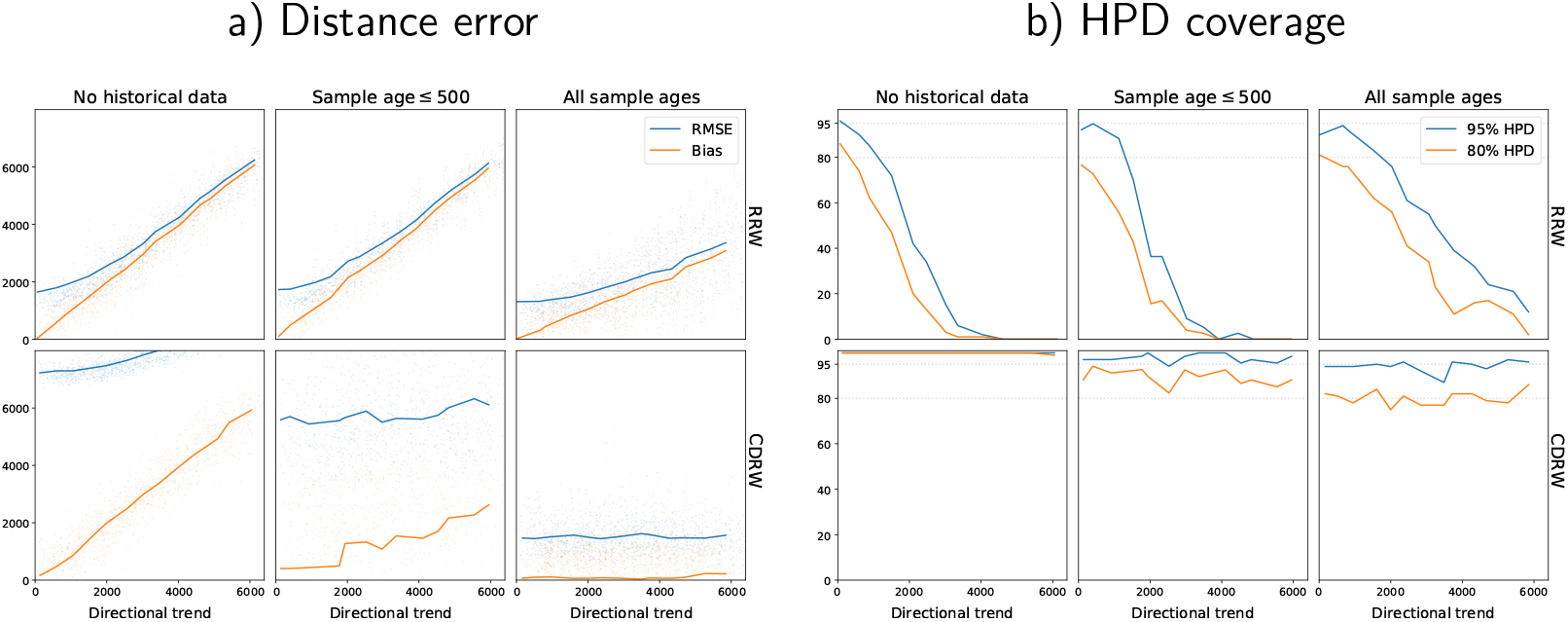
The performance of phylogeographic reconstructions of the root based on the MigSim simulations with varying levels of observed trend. **a)** The RMSE (blue) and the bias (orange) of the reconstruction. The dots represent single simulation runs, the lines interpolate between the average results for a specific setting for *μ* (trend). **b)** The empirical coverage of the 80% (blue) and 95% (orange) credible regions.

**Figure 3:**
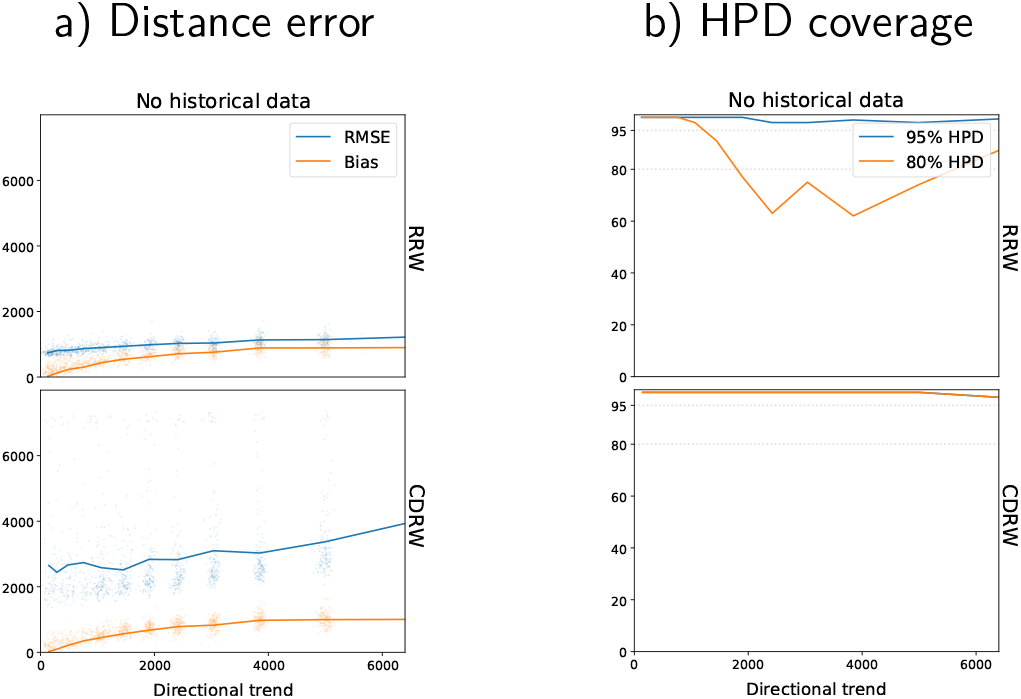
The performance of phylogeographic reconstructions of the root based on the ExpSim simulations with varying levels of observed trend. **a)** The RMSE (blue) and the bias (orange) of the reconstruction. The dots represent single simulation runs, the lines are averages across all runs with a specific setting for *α* (sector angle). b) The empirical coverage of the 80% (blue) and 95% (orange) credible regions.

In the absence of historical data, directed migrations lead to significant errors in the phylogeographic reconstruction. This can be seen in Figure 2a in the leftmost column (no historical data): The reconstruction bias increases linearly with the simulated directional trend. If we observe a directional trend of 5000 km, the reconstruction is off by about 5000 km too. This holds for reconstructions using the RRW (top row) as well as the CDRW (bottom row) model. This is expected, since the CDRW model needs historical data to be calibrated [22]. The RMSE (blue line) of the CDRW reconstruction is even significantly higher, due to a higher variance in the posterior. A look at the HPD coverage statistic shows that this increased variance is an effect of a very agnostic or even underconfident model, where the 95% and even the 80% HPD regions always cover the simulated root. In the RRW model, in contrast, the corresponding HPD coverage approaches 0 with increasing directional trends, as expected in the case of an increasingly significant mismatch between the model assumptions and the data.

Including historical data in the analysis reduces the error in both the RRW and the CDRW model. If we include sampled historical locations from the whole time period, the CDRW model is able to estimate the directional trend from the data, which leads to a bias close to 0 and 80%/95% HPD coverage around the ideally expected values of 80%/95%, respectively (bottom row, right). The reconstruction of the RRW model on the other hand only flattens the slope at which the error increases. At an observed trend of 5000 km the reconstruction bias is still above 2500 km. Furthermore, the model is still overconfident, visible in the drop of the HPD coverage. Finally, we find that including historical locations at shallow time depths of up to 500 years does not significantly improve the reconstruction with either model.

In expansion scenarios (ExpSim) we see a very different picture. Even in the absence of any historical information the reconstruction error levels off far below what we would expect in the MigSim scenario, as can be seen in the top panel of Figure 3a. After an initial increase of the reconstruction error, even at an observed directional trend of 7000 km the reconstruction bias does not exceed 1000 km. Since the exact values of these errors (bias and RMSE) are influenced by the scaling parameter (kilometers per cell) and the shape of the simulated constraints, it is the more general results that we want to emphasize: a) The bias does not increase linearly with the observed trend, but clearly stagnates and b) the estimated HPD regions cover the true homeland very consistently (Fig. 3b). These results turn out to be independent of the phylogeographic model used. Resorting to an explicit directional model (CDRW or RDRW) only increases the RMSE, through a higher variance in the posterior distribution.

Furthermore, we show in a sensitivity analysis (supplementary material 4) that the tree size does not notably affect the reconstruction quality (5) and that our general findings for the ExpSim scenario still hold if areas are allowed to overlap (5.1).

## 5 Discussion

In summary, the results of the simulation study reveal that the reconstruction quality is vastly different in different movement scenarios:

- The migration scenarios (MigSim) lead to a severe bias in the reconstruction that grows proportionally with the directional trend in the migration. This bias remains when including recent historical samples and is independent of the tree sizes (see supplementary material 5).
- In expansion scenarios (ExpSim), the model shows some uncertainty about the precise location of the origin, but no severe reconstruction errors, even under strong directional trends. The successful reconstruction does not rely on historical samples, is robust to overlapping language areas and changes in the tree size (see supplementary material 5 and 5.1).

In what follows, we discuss why phylogeographic methods perform so much better in the ExpSim scenario (5.1) and we make recommendations for future work (5.2).

### 5.1 Differences in reconstructability

Why are we able to reconstruct the origin in the expansion scenario, while the same seems to be impossible for the migration scenario? We visualize the differences in Figure 4 and show statistics characterizing the different scenarios in Figure 5. In a typical migration scenario (Fig. 4, left column), the diversification and the movement processes are independent. If there are no other factors causing variation in the birth or death rates, we expect a balanced tree topology. The migrations follow a random walk along this tree with a bias in one direction, as visualized in the middle row of Figure 4. Without any historical information, there is no way for a reconstruction algorithm to identify this direction. The most sensible (parsimonious) reconstruction is achieved by a homeland at the center of mass of the recent locations (Fig. 4 bottom left).

**Figure 4:**
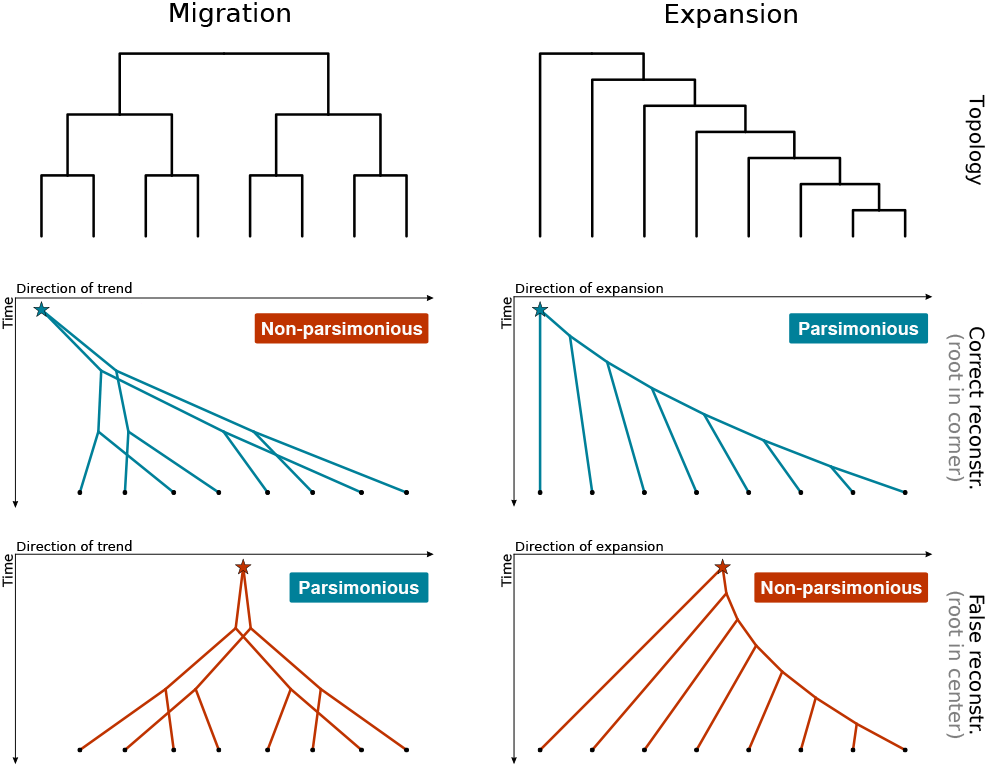
Visualizations of two scenarios of directional trend: migration (MigSim) and expansion (ExpSim). In all plots the y-axis represents time. The top row depicts typical tree topologies (balanced vs. imbalanced) assuming no further variation in birth and death rates. The middle row shows typical migration patterns in one spatial dimension (parallel migrations vs. nested splitting and expansion). The bottom row shows how a hypothetical reconstruction of the homeland at the center of gravity would lead to more vs. less parsimonious reconstructions. In the expansion scenario the more parsimonious case coincides with the correct reconstruction.

**Figure 5:**
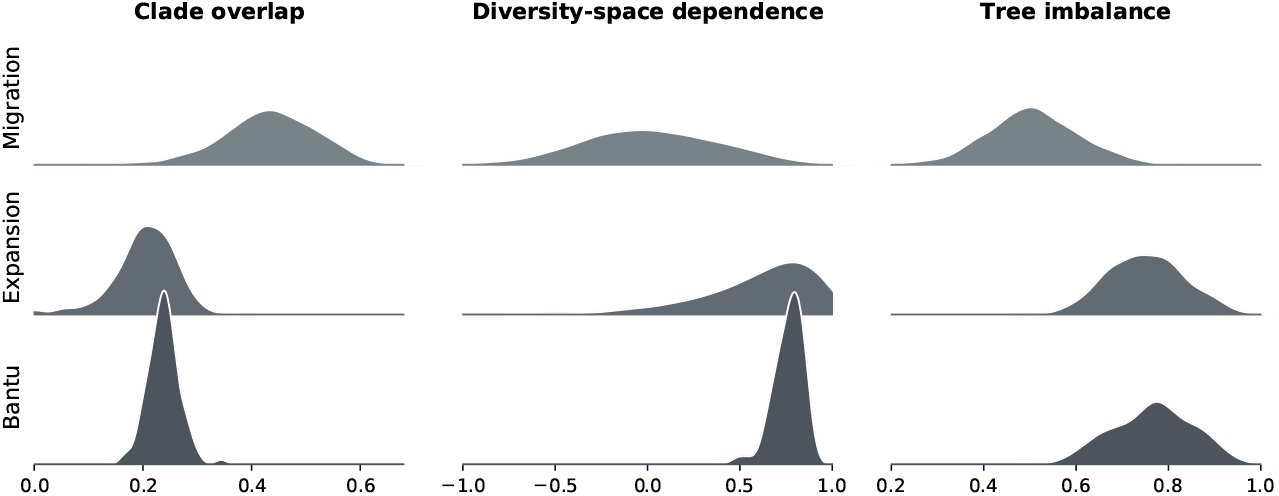
Three descriptive statistics 1) clade overlap 2) diversity-space dependence and 3) tree imbalance (see supplementary material 5) evaluated on samples from the MigSim simulation, the ExpSim simulation, and the posterior distribution of the Bantu phylogeny.

Expansions lead to different spatial and phylogenetic patterns compared to migrations, even if they show the same level of observed directional trend (right column of Fig. 4). First, expansions do not cause all languages to move in one direction. Actually, every language tends to stay stationary and it is only the diversification and growth into new areas that is forced to proceed in one direction due to geographic constraints. In turn, clades which stay stationary are more limited in their space for further expansion and diversification. As a result the region of the homeland will mostly be populated by clades that stopped to diversify and to migrate a long time ago (compared to the languages at the frontier of the expansion). In Figure 4 (right column) we illustrate what such an expansion typically looks like, with an imbalanced tree topology, where languages split off one by one. The languages splitting off earlier remain closer to the homeland, while the expansion continues to spread, causing the directional trend (Fig. 4, middle right). Figure 4 (bottom right) shows that a reconstruction of the true directed expansion is actually more parsimonious than a spread from a hypothetical central homeland, which would imply multiple parallel migrations of early-split languages towards the left.

We propose three descriptive statistics to characterize the different patterns seen in the MigSim and ExpSim scenarios (see supplementary material 5 for details): The **clade overlap** score measures how much languages from different clades overlap in space. Under the ExpSim scenario clades are tightly connected, since the areas are generally stationary and cannot cross each other (this constraint is relaxed in supplementary material 5.1; overlap score of 0.20±0.06). In the MigSim scenario languages can move and cross each other freely, leading to an increased clade overlap (0.43±0.09).

The **diversity-space dependence** captures the fact that diversification in the ExpSim scenario is directly governed by the area of a language, while the two processes of spread and diversification are independent in the MigSim model. This leads to a dependence score of 0.60±0.34 for ExpSim and 0.02±0.36 for MigSim.

Finally, the **tree imbalance** score is high in the ExpSim scenario (0.75±0.09), because the growth of earlier clades is inhibited through geographic constraints and other clades (as described above and visualized in Figure 4). In the MigSim scenario the tree shape follows a common birth-death process, leading to a relatively balanced tree topology (0.50±0.10).

These descriptive statistics may help to identify the spread of a language family as a migration or an expansion scenario, which in turn indicates whether we can trust the phylogeographic reconstruction. Beyond this, we also want to highlight a basic property of phylogeographic reconstructions (using the RRW model): The reconstructed root will always fall within the convex hull of the sampled locations. As shown in [33], the mean location of the internal nodes can be expressed as a weighted average of the sampled locations, directly implying that they must be within the bounds of the convex hull. Clearly, phylogeography is not an adequate reconstruction tool if a migration from outside the sampled area (or a loss of the languages in the area of the homeland) is a plausible hypothesis.

### 5.2 Implications for linguistic phylogeography

In the remainder of this section we explore the application possibilities and restrictions of current phylogeographic methods, and make recommendations for judging the adequacy of phylogeographic models to reconstruct the spatial evolution of languages. The simulation study presented in this paper shows that phylogeography provides faithful reconstructions of expansions, but fails severely on directed migrations. To translate these findings into practical decisions for the reconstruction of a real language family, researchers need to judge whether the history of those languages suggests an expansion or a migration scenario. If a plausible hypothesis about the history of the languages involves directed migrations, phylogeography is not an appropriate tool for reconstruction or hypothesis testing.

We now discuss the Bantu languages as an example of a linguistic expansion with an existing phylogeographic analysis (Grollemund et al. [12], available as part of DPLACE [41]). The spread of the Bantu languages goes hand in hand with one of the major expansions of agriculture on the planet. As the farming/language dispersal hypothesis posits [19], the availability of agriculture explains a continuous spread of the Bantu people and languages into most of southern Africa. The geographical constraints of this dispersal are roughly defined by oceans, deserts (Sahara, Kalahari), and land occupied by related Niger-Congo societies (West Africa). In fact, the Bantu languages nowadays extend over most of the geographic area bounded by these constraints. The phylogeny of the Bantu languages exhibits similar patterns to those of the expansion (ExpSim), and not those of a migration scenario (MigSim; Figure 5): The phylogenetic tree is imbalanced (imbalance score of 0.77±0.08), with little clade overlap (overlap score of 0.24±0.03) and the clades that split off and became stationary also diversified less than the rest of the tree, which was able to expand into a wider territory (leading to a diversity-space dependence score of 0.76±0.08). This process is quite clearly visible in the reconstructed expansion (Figure 6). All these indications would support the notion that the spread of the Bantu languages is well described as an expansion scenario and, indeed, the phylogeographic reconstruction in [12] very accurately matches the previously proposed homeland in the Grassfields region in Cameroon (supported by strong scientific consensus, based on archaeological [42, 43], linguistic [44], and genetic evidence [45]). Similar patterns where evidence is suggestive of multiple nested splits corresponding to chained movements into new territories are found all over the globe. For example, the Austronesian expansion is also characterized by an imbalanced tree topology and multiple successive waves of expansions towards the east [46]. In the Austronesian case, geographical constraints, coupled with the development of seafaring technology seem to have determined the pace and direction of the expansion [6].

**Figure 6:**
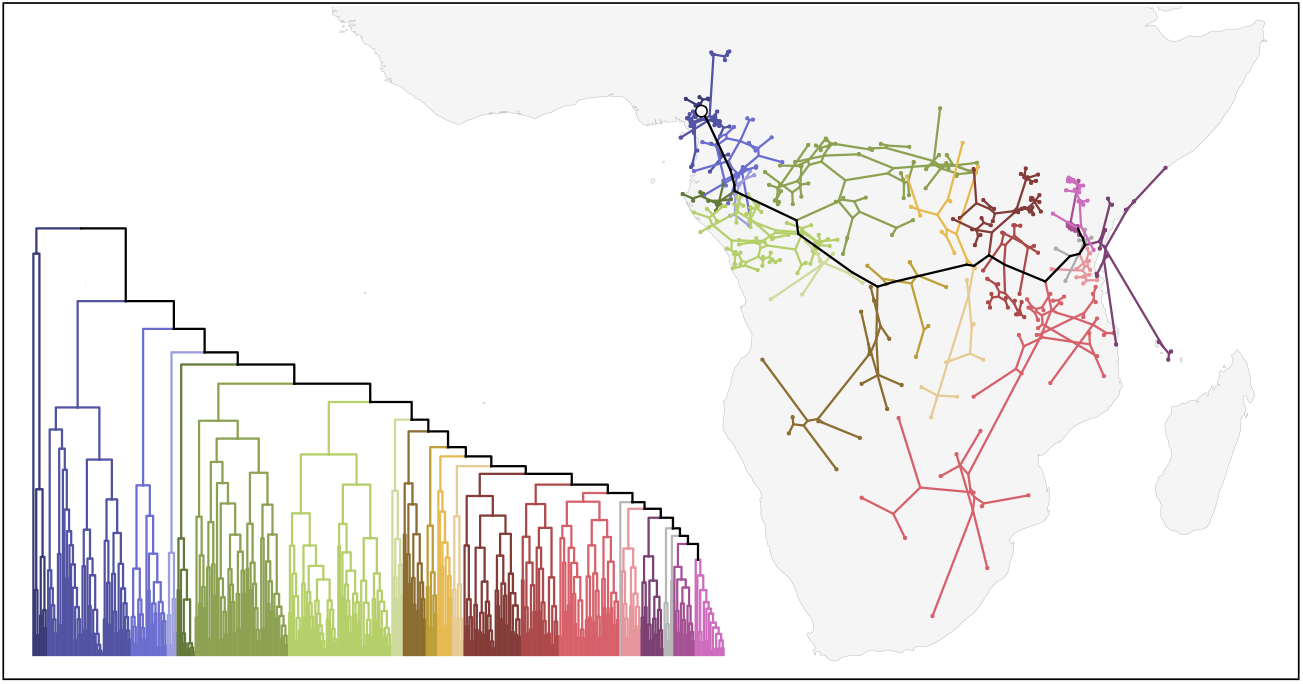
The phylogeny and spatial spread of the Bantu languages according to [12]. The colors mark the clades, splitting off one after another from the backbone of the expansion.

If the history of the languages under study does not align with the expansion scenario, we cannot warrant the same optimism expressed above. If the history involved directed migrations we would even expect the reconstruction to fail. This includes cases of demic migrations as well as scenarios where languages were replaced and pushed out of their initial territory in a language shift. The latter is clearly observable in the case of the Celtic languages, which once extended across vast areas of Europe. But since the population in these areas shifted to Germanic and Romance languages, Celtic is now only spoken in north-western France (Brittany) and on the British Isles, making a phylogeographic reconstruction of their homeland impossible (as can be seen in [11]).

Finally, we want to note that even in an expansion scenario we have to be aware of the opacity of history. The Bantu expansion beautifully illustrates how the first clades inform the reconstruction of the homeland despite a strong directional trend in the later expansion. But what if these first clades were missing or migrated away? In the supplementary material (S6The effect of missing first cladesfigure.6) we show how the reconstruction of the Bantu homeland would fail dramatically in this hypothetical scenario. This emphasizes the importance of complete sampling, especially in early splits and in the regions around a hypothesized homeland. Furthermore, including an outgroup of languages not part of the family under study will stabilize the reconstruction of the homeland. Specifically, [12] included an outgroup of Jarawan and Grassfields languages with the Narrow Bantu languages.

Besides the findings in this study, a reconstruction has to be checked against the historical context. Clearly, including records of ancient languages will improve the reconstruction, but this is often not possible. Instead, therefore, the reconstruction can be corroborated by archaeological or genetic findings. Especially findings of cultural artifacts and practices associated with the language family and knowledge about the spread and development of domesticated crops can give a sense for the plausibility of the reconstruction. More generally, qualitative discussions, as is common in historical linguistics, are an important complement to quantitative, phylogeographic reconstructions. For example, the embedding in a deeper historical context (such as the spatial distribution of other Niger-Congo languages for Bantu), knowledge about contacts with other language families (such as contacts between Indo-European and Uralic languages), or knowledge about trade routes (such as the river-based Arawakan expansion discussed in [47]), as well as detailed studies of reconstructed vocabulary items (as e.g. in [48] for Mayan and [49] for Uto-Aztecan) can be taken into consideration to further support a geographical reconstruction.

## 6 Conclusion

Directional trends in the spatial evolution of languages are a clear case of model misspecification for the most widely used models in phylogeographic analyses. We have demonstrated the effect of directional trends on the reconstruction of language spread in two movement scenarios-migrations and expansions-with corresponding historical interpretations. The effects on the reconstructions differ greatly between the two scenarios: migrations (e.g. Chadic and Celtic) make a correct reconstruction impossible, while expansions (e.g. Bantu and Austronesian) only lead to minor imprecisions in the reconstruction, even under severe directional trends. The message to researchers applying phylogeographic methods accordingly depends on the historical scenarios they are investigating: in scenarios of continuous expansions, where a potential directional trend was the result of geographic constraints, our findings support the validity of phylogeographic analysis. In cases where external changes, such as environmental factors or replacement by other languages, might have caused a displacement to the currently inhabited areas, our results strongly discourage the use of phylogeographic reconstruction methods.

## Data accessibility

All Python scripts and instructions for reproducing our results can be found in the following repository: https://github.com/NicoNeureiter/drifting_into_nowhere/.

## Author’s contributions

N.N. and P.R. designed the research; N.N. implemented the simulations and performed the experiments; B.B. and R.v.G. interpreted the results in a linguistic and historical context; R.W. coordinated the study; N.N. and P.R. wrote the paper with contributions from R.v.G., B.B., R.W. All authors gave final approval for publication and agree to be held accountable for the work performed therein.

## Competing interests

We declare we have no competing interests.

## Funding

This research was supported by the University of Zurich Research Priority Program ‘Language and Space’, the Swiss National Science Foundation Sinergia grant no. CRSII3_160739; ‘Linguistic Morphology in Time and Space’ (LiMiTS), the European Research Council, ERC Consolidator grant no. 818854 ‘South American Population History Revisited’ (SAPPHIRE) and the NCCR Evolving Language.

## Acknowledgments

We would like to thank Curdin Derungs, Péter Jeszenszky and Nour Efrat-Kowalsky for their valuable input, discussions and feedback.

## Notes

https://github.com/NicoNeureiter/drifting_into_nowhere/

### Competing Interest Statement

The authors have declared no competing interest.

