## Supporting Information for "Can Bayesian phylogeography reconstruct migrations and expansions in human history?"

#### Contents

**S1 Simulations** **S1**
S1.1 Migration simulations (MigSim) . . . . . S1
S1.2 Expansion simulations (ExpSim) . . . . . S2

**S2 Reconstruction** **S2**

**S3 Evaluation** **S3**

**S4 Sensitivity analysis** **S4**
S4.1 Sensitivity: Tree size . . . . . S4
S4.2 Sensitivity: ExpSim area overlap . . . . . S5

**S5 Descriptive statistics** **S6**
S5.1 Clade overlap . . . . . S6
S5.2 Diversity-space dependence . . . . . S8
S5.3 Tree imbalance . . . . . S8
S5.4 Migrations, expansions, and Bantu . . . . . S8

**S6 Effect of missing first clades** **S10**

This document provides supporting information for the main article "Can Bayesian phylogeography reconstruct migrations and expansions in human history?". It includes a detailed description of each step in the simulation study (S1-S3), a sensitivity analysis (S4), a detailed explanation of the descriptive statistics used in the discussion of the main paper (S5) and a figure, illustrating the effect of missing first clades on the reconstruction (S6).

### S1 Simulations

We simulate the evolution of phylogenetically related languages in space as two temporal processes: Movement and diversification. That is, the simulated languages move in space and randomly split to form descendant languages, which then continue to move and diversify themselves. While diversification is important to simulate phylogenetically related languages, our focus lies on the movement process. We introduce two simulation scenarios corresponding to the two historical processes of migration and expansion.

#### S1.1 Migration simulations (MigSim)

We model migrations as directional random walks, where languages are represented by points in space. These points move stochastically with a bias in one direction. The direction and strength of this bias is controlled by the parameter $\mu$. Examples for different levels of $\mu$ are displayed in Figure S1. Other parameters of the migration simulations are $T$, the total time span of the simulation (from the first split to the current time); $N$ the expected number of leaves (i.e. sampled languages) in the tree; and $\sigma$, the expected distance covered due to undirected movements over the whole expansion period. Here, we set these parameters to values that seem realistic for the expansion of a language family ($T = 5000$ years, $N = 100$ nodes and $\sigma = 2000$ km). However, we want to emphasize that the exact values do not matter for our findings. In our results we show more generally how the reconstruction quality changes when increasing $\mu$, for a fixed $\sigma$. A sensitivity analysis on the number of nodes shows that varying the tree size does not change our findings (see S4).

We implement directional random walks in discrete time steps of duration $\Delta_t$ (in our experiments set to 1 year). In every time step each language makes a random move according to a Gaussian distribution:

$$X_{t+\Delta_t} \sim \mathcal{N}\big(X_t + \mu_{\text{step}},\ \sigma^2_{\text{step}}\big) \tag{1}$$

The free parameters in this process are the step mean $\mu_{\text{step}}$ and the step variance $\sigma^2_{\text{step}}$. Since these steps are arbitrary units of our simulation, we aggregate them into more meaningful quantities: the total bias $\mu$ and total standard deviation (or total expected diffusion distance) $\sigma$, defined as

$$\mu = T \cdot \mu_{\text{step}} \tag{2}$$

$$\sigma = \sqrt{T} \cdot \sigma_{\text{step}} \tag{3}$$

This gives rise to a reformulated step distribution

$$X_{t+\Delta_t} \sim \mathcal{N}\big(X_t + \mu/T,\ \sigma^2/T\big) \tag{4}$$

At the same time each language has a certain probability to split into two new languages, which then continue to perform independent random walks. In order to simulate historical data from extinct languages (fossils), each language has a certain probability to go extinct. This is a common birth-death process, which is controlled by the birth rate $\lambda$ and death rate $\nu$. We set $\nu = \frac{\ln N}{4T} = 0.00023$ and $\lambda = 5\nu = 0.00115$, which after $T = 5000$ years results in an expected number of languages of $N = 100$. To ensure comparability between the scenarios, we remove outliers from the simulation results. We consider a result as an outlier if the number of extant languages is too small or too large (below 40 or above 200).

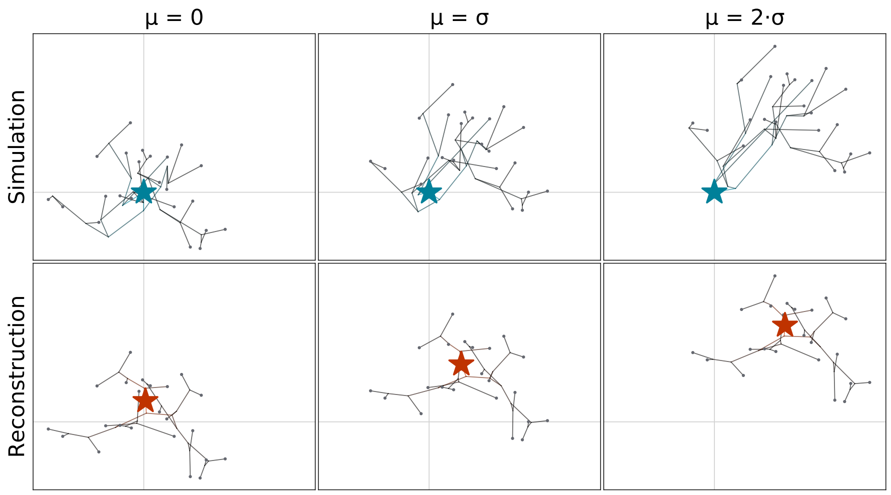


Figure S1: Examples of three simulations and corresponding reconstructions. The top row shows the simulated trees plotted in space with the root $(0, 0)$ marked by a blue star. In the bottom row the reconstructed tree can be seen with the root in red. The columns represent different levels of directional trend in the simulation. The trend increases from left to right ($\mu = 0$, $\sigma$, $2\sigma$).

#### S1.2 Expansion simulations (ExpSim)

We propose expansion simulations, which can also be subject to directional trends, however, not through an inherent directional bias, but through geographic constraints, forcing an expansion in one direction. Geographic constraints could be barriers, such as mountains or oceans, which are difficult to traverse, or simply disfavored regions, where for example an important crop does not grow. We capture this constrained expansion scenario in a simulation based on the following ideas (Figure S2): Languages are represented by areas consisting of cells of a grid. These areas randomly expand over time into free neighboring cells, i.e. cells that are not occupied by another language or blocked by geographical constraints. In a sensitivity analysis (see S4) we relax this assumption and allow up to three languages to occupy the same cell. With increasing area a language becomes more likely to split into two new languages, which in turn continue to expand separately.

As illustrated in Figure S2, the ExpSim simulation is defined by two parallel processes: Growing and splitting of areas. Growing is controlled by a parameter $p_{\text{grow}}$. At every step of the simulation an area adds a random cell from its neighborhood with probability $p_{\text{grow}}$. At a certain randomly chosen size a language splits up into two. In our experiments we choose the split-size uniformly at random and for every language independently between 70 and 100 cells. Again, the exact values do not significantly influence the results. We chose the distribution of the split-size to roughly match the expected tree size in the MigSim experiments. In order to make space and time comparable we define a step to correspond to one year and a cell to correspond to $100 \times 100$ kilometers.

To introduce a directional trend into the simulation, we model geographical constraints. In reality these constraints (e.g. deserts, mountains or seas) may have arbitrary shapes, but to systematically investigate the properties of constrained expansions we confine the simulations to geometric examples: The first language starts to expand in the corner of a circular sector of angle $\alpha$ and is only allowed to grow within this sector. Leaving the sector completely open ($\alpha = 2\pi$) allows a concentric expansion in all directions. Choosing the sector very tightly ($\alpha \to 0$), forces the expansion to move in one direction out along this narrow sector. In this way we can control $\alpha$ to achieve different levels of directional trend in the expansion. To compare this trend to the inherent bias parameter in the MigSim scenario, we will introduce a measure for the observed trend in Subsection S3.

### S2 Reconstruction

To evaluate the performance of phylogeography, we start from a setting which is optimistic compared to real-life applications: We fix the phylogenetic tree to the true (simulated) one and perform

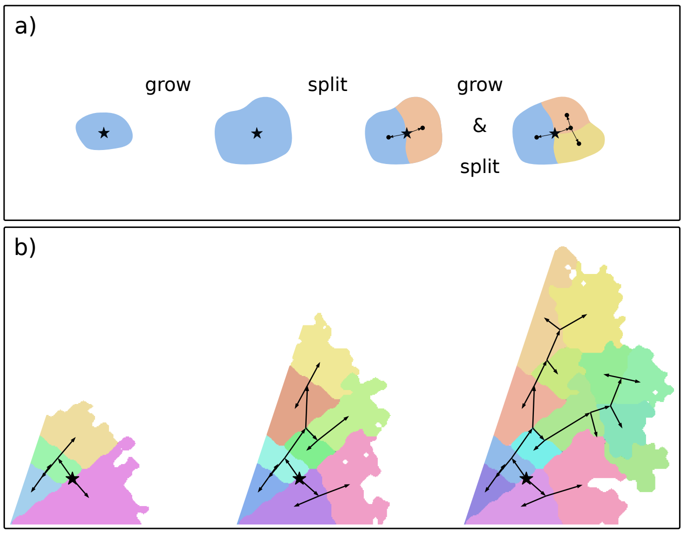


Figure S2: Two illustrations of the expansion simulation: a) Every language is represented by a coloured area, which over time grows into free surrounding areas and splits up into new languages. The result is a phylogenetic and geographic expansion, as we can further see in b): The expansion started at the black star and grows into new free space (visualized at three time points). Free space in this case is only available in a 72° sector. The phylogenetic tree is visualized by the black edges, leading from the root (black star) to the different areas, representing the extant languages.

phylogeographic analysis based on this tree and the tip locations. In a real case study the tree would have to be reconstructed as well. Fixing the tree is a valid simplification of the experiment, because we are really only interested in the evaluation of the geographic reconstruction. The result of the analysis is a reconstruction of the locations at the internal nodes of the tree, in particular the root location (the homeland). We perform this phylogeographic analysis in BEAST 1.10.4 [1], where the original RRW model [2] and extensions to directional random walks [3] are implemented.

For the phylogeographic analysis we test the relaxed random walk (RRW) and the two directional random walk models CDRW and RDRW. All models were set up with variable rates according to a log-normal clock. The prior settings of the analysis are in line with commonly used values in previous analyses (e.g. [4]). Moderate changes in the prior on the diffusion rate and the variance of the log-normal clock did not show any notable effects. In CDRW and RDRW the prior on the drift parameter (directional bias) heavily influences the reconstruction in cases where we do not provide enough fossils to calibrate the parameter. The effect of the prior ranges from a fall-back to the RRW model, for priors that are very narrow around 0, to a highly variant reconstruction that would allow for directed migrations from any point on the map, for very wide priors.

#### S3 Evaluation

A Bayesian phylogeographic reconstruction, as the one described above, provides a posterior distribution over possible root locations. We are interested in two properties of this reconstruction: 1) How far do we expect the reconstructed location to be from the true homeland? 2) How certain is the model about its reconstruction? We measure these two properties using the following metrics:

1. We can measure the Euclidean distance between every location in the posterior distribution and the true root to see how far the reconstruction is off. Doing this per posterior sample and taking the root of the mean squared error gives the common RMSE metric:

$$\text{RMSE} = \sqrt{\sum_{m=1}^{M} \left\| X - \hat{X}_m \right\|^2 / M} \tag{5}$$

   Here $X$ is the true location of the homeland and $\hat{X}_m$ are the reconstructed locations, taken from all posterior samples over multiple simulation runs. The RMSE is composed of systematic errors (bias) and errors due to variance (in the simulation and in the reconstruction).

Since we are mainly interested in systematic errors arising from directed migrations we also present the bias of our reconstructions:

$$\text{bias} = \left\| X - \sqrt{\textstyle\sum_{m=1}^{M} \hat{X}_m / M} \right\| \tag{6}$$

As an illustration of the behavior of these two metrics consider the examples shown in Figure S1. With growing directional trend $\mu$, the reconstructed root (red star) will be located further away from the simulated one (blue star), causing both the RMSE and the bias to grow. The difference between the two scores would mostly show in scenarios with low directional trend ($\mu$ close to 0), where the bias would approach 0, while the RMSE still shows errors due to variance.

2. We measure the Bayesian highest posterior density (HPD) region coverage of the root location for different HPD thresholds (80% and 95%). This entirely ignores the amplitude of the error, but shows whether the uncertainty expressed by the posterior reflects the observed error. If so, the HPD coverage will match the corresponding HPD threshold, i.e. the 95% HPD region should cover the root in 95% of the simulations.

In order to compare these evaluation metrics between the different scenarios, we want to measure directional bias in a unified way. In the case of migrations, the directional bias is introduced through an inherent trend parameter $\mu$, in the expansion simulations it is controlled by the sector angle $\alpha$ (and in a real case study we usually do not know the factors driving the directed migrations at all). To compare these scenarios we measure the **observed trend** $\hat{\mu}$, which we defined as the distance between the homeland of an expansion and the mean of the final locations (at presence):

$$\hat{\mu} := \left\| \textstyle\sum_{n=1}^{N} x_n / N - X \right\| \tag{7}$$

Here $X$ represents the root of the expansion and $x_n$ are the tip locations (contemporary languages). We will use this statistic to compare the reconstruction error of different simulations showing the same observed trend.

### S4 Sensitivity analysis

In this paper we have presented two simulation scenarios: migration simulations (MigSim) and expansion simulations (ExpSim). These two scenarios lead to very different performance in the reconstruction of the simulated histories, which is a core part of our findings. To ensure that these findings are really stable properties of the two simulation scenarios and do not depend on the specific settings of some parameters, we performed a sensitivity analysis, which we present here. In the first analysis we will test the effect of the size of the simulated phylogenetic trees (S4.1). In the second analysis we explore the effect of relaxing the assumption that languages can not overlap in the expansion simulations (S4.2).

#### S4.1 Sensitivity: Tree size

In the MigSim scenario it is fairly straightforward to vary the tree size, while keeping other parameters fixed: We simply set the birth-rate to $\lambda \in [0.0009, 0.0012, 0.00143]$ (and the death-rate accordingly to $\nu \in [0.00017, 0.00023, 0.00029]$), leading to an expected tree size of 50, 100 and 300, respectively. The results of these experiments are shown in Figure S3. We can see that the results are, indeed, very robust to variations of the tree size.

For the expansion simulations there is no analytical fomula to compute the expected tree size from the parameters, thus adjusting the tree size requires some tuning. As in the MigSim scenario, we change the diversification process, in this case by adjusting the size at which languages split. In the default case the split size was randomly chosen between 70 and 100, leading to a tree size of about $N = 100$. We varied this split size range between $(140, 200)$ and $(25, 33)$ to obtain trees with about 50 and 300 leaves, correspondingly. As we can see in Figure S4, changing this parameter has a more notable effect here. Firstly, the simulations show a slightly lower observed trend with smaller trees (i.e. when we increase the split size of the areas). Secondly, the reconstruction error increases

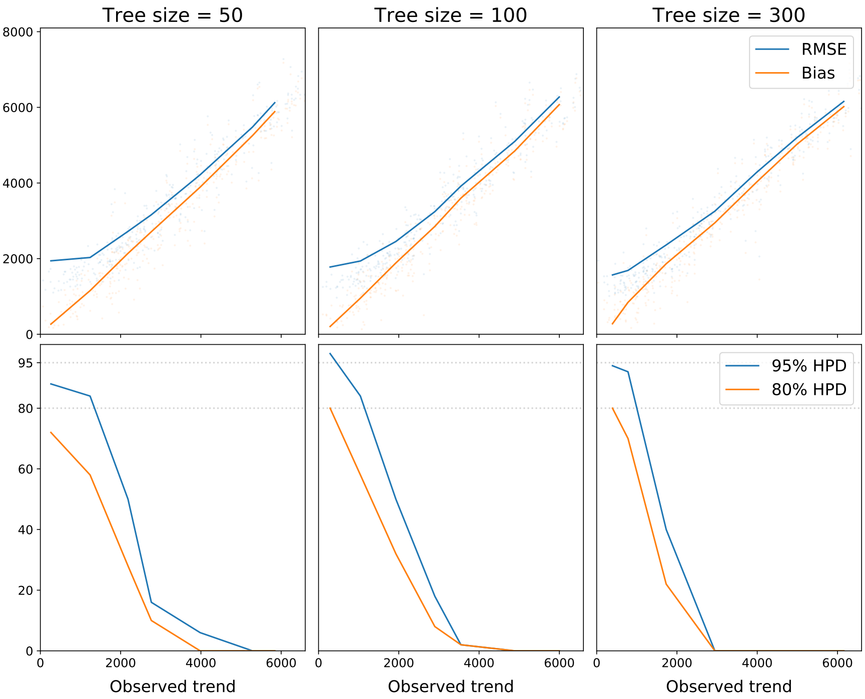


Figure S3: Results for the MigSim scenario, with varying tree size. The top row depicts the bias and RMSE of the reconstruction, the bottom row the 80% and 95% HPD coverage.

and the HPD coverage decreases for small trees. Despite these differences our main findings hold: The reconstrucion error (measured as RMSE and bias) levels off and is far away from the linear increase seen in the MigSim scenario. While the coverage does fall below the desired values of 80% and 95%, respectively, for small trees with certain trend values, in almost all settings the confidence regions seem to overestimate the uncertainty (i.e. we can trust the confidence regions provided by the reconstruction).

#### S4.2 Sensitivity: ExpSim area overlap

In human history multilingualism and coexistence of different populations in the same place are the rule rather than the exception. In contrast to this, the original version of the ExpSim simulation only allows language areas to grow into free cells (not inhabited by any other population). The motivation for this is the scenario of populations expanding into new areas, where their languages where not spoken before, e.g. in search of available land to grow their crop. Still, we want to be sure that our findings do not break down if this hard constraint is relaxed. To do this we ran an analysis on a modified simulation where overlaps between areas are allowed within two constraints:

1. The number of languages per cell must not exceed 3. Without a limit on the number of languages per cell, the languages would diversify exponentially within an area before they even get to expand into new areas (which happens at a quadratic rate). Simulating a significant geographic expansion without this constraint would thus lead to unrealistic and computationally problematic clusters of many languages in the same area.

2. Languages are more likely to expand into free cells than occupied cells. The probability of growing into an occupied cell is reduced by a factor of $\rho_{\text{overlap}}$ which we vary between 0 and 1, where 0 corresponds to the standard setting with disjoint areas and 1 corresponds to a setting where the occupancy of a cell does not matter (up to the limit of the first constraint, i.e. 3 languages).

As can be seen in Figure S5 the reconstruction bias increases when we allow language areas to grow into each other, but even in the extreme scenario of $\rho_{\text{overlap}} = 1$, where previous (up to three) languages are completely ignored, the error levels off. That is, our findings regarding the ExpSim scenario indeed withstand the relaxation of the overlap constraint.

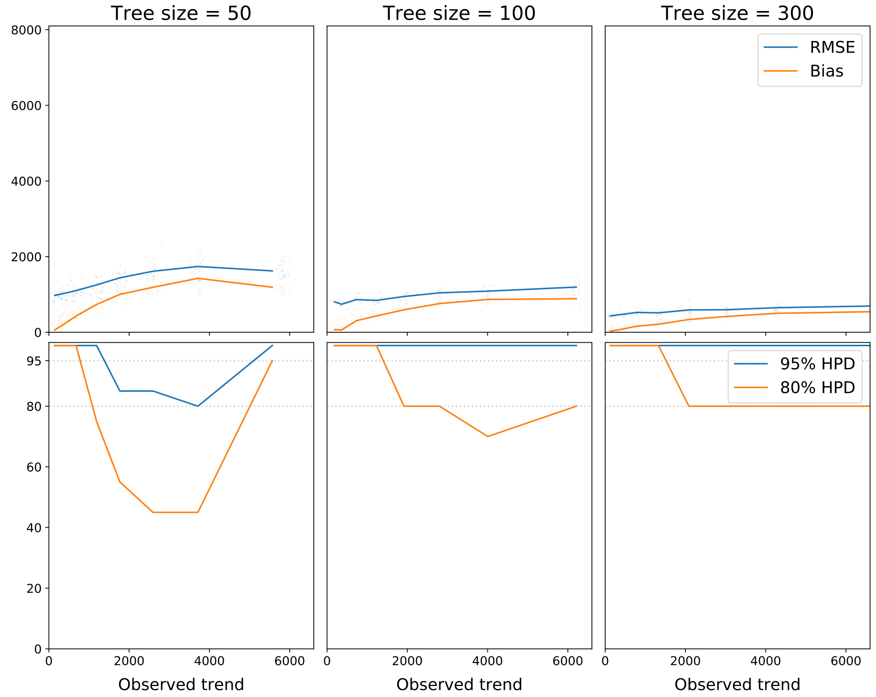


Figure S4: Results for the ExpSim scenario, with varying tree size. The top row shows the bias and RMSE of the reconstruction, the bottom row the 80% and 95% HPD coverage.

### S5 Descriptive statistics

The two simulation scenarios for migrations (MigSim) and expansions (ExpSim), show very different patterns, not only in the spatial distribution of the languages, but also in the tree shape. We visualize these differences in Figure 4 of the main paper.

In the MigSim scenario, the diversification and the movement processes are independent, and under common tree priors we would expect a relatively balanced tree topology. The movement follows a random walk along this tree with a bias in one direction, as visualized in the middle row of Figure 4. In the expansion scenario on the other hand, the diversification and spatial expansion are strongly dependent. This leads to different spatial and phylogenetic patterns from random walks (right column of Figure 4).

In the ExpSim scenario there is no intrinsic bias for languages to move in one direction or to move at all. Actually, every language tends to stay sedentary and it is only the diversification and spatial expansion that is forced to progress in one direction due to geographic constraints (and closer areas being occupied by previous sedentary languages). As a result the region of the homeland will always be populated, mostly with languages that stopped to diversify and to migrate a long time ago (compared to the languages at the frontier of the expansion). In Figure 4 (right column) we illustrate what such an expansion typically looks like, with an unbalanced tree topology (top), where languages split off one after another. The languages splitting off earlier remain closer to the homeland, while the expansion spreads away, causing the directional trend (middle row of Fig. 4).

The observations made above regarding these different processes can be expressed by three measurable properties, which we describe below. The spatial dimensions of these properties are based on point locations of languages. In the ExpSim scenario we take the centroid of an area as the point location.

#### S5.1 Clade overlap

The ExpSim scenario enforces disjoint language areas, which cannot cross each other in the course of the expansion. This means that the clades of the phylogeny will cover non-overlapping geographic areas. In the MigSim scenario on the other hand, the languages can move freely and independently

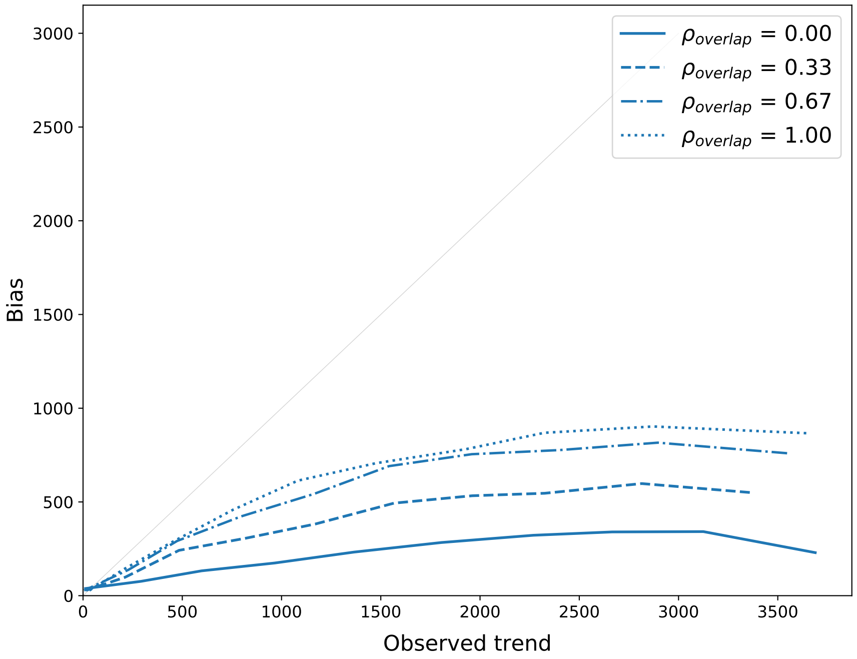


Figure S5: The reconstruction bias (see Section S3) in the ExpSim scenario with varying overlap coefficient $\rho_{\text{overlap}}$.

in space, leading to significant overlap between the clades. To quantify the concept of clade overlap in a measurable score, we need to do two things:

1. Define the clades to be compared.
2. Measure the overlap between the clades.

**Clades** For clades to be comparable we fix a tree height $h$ and define clades as all the sub-trees starting in lineages at that height. We (somewhat arbitrarily) set $h$ to half the total tree height. The rationale here is that we want enough clades to compare, but also clades of a sufficient size to measure a meaningful overlap (i.e. balancing out the number of clades and clade size).

**Join count statistic** To quantify the overlap between clades, we follow a simple observation: If clades overlap, nodes will have spatial neighbours from different clades, if they do not overlap, the spatial neighbours will mostly be from the same clade. We use the Delaunay triangulation to define spatial neighbourhood (so-called 'natural neighbors') and measure the fraction of neighbours in the same clade according to the *join count statistic*. More specifically, we count all edges $(i, j)$ in the Delaunay graph $G$ where both nodes are in a certain clade $C \subseteq V(G)$ and denote the result by

$$n_{\text{bb}}(G, C) \coloneqq |\{(i, j) \in G | i \in C \wedge j \in C\}|.$$

Clearly, $n_{\text{bb}}$ strongly depends on the size of the clade (the larger $C$, the higher we expect $n_{\text{bb}}(G, C)$ to be). Thus, we define a normalized *compactness* score $\alpha_C$, such that $\alpha_C = 0$ corresponds to a scenario where clades are completely randomly mixed in space (i.e. no spatial auto-correlation of clades), while $\alpha_C = 1$ corresponds to a scenario of maximal connectedness within a clade. We perform this normalization by computing the expected join count in the former and the latter scenario (denoted by $n_0$ and $n_1$). We can compute $n_0$ by shuffling the clade labels of all locations and compute the $n_{\text{bb}}$ score based on the shuffled labels:

$$n_0(G, C) = \mathbf{E}_\pi[|\{(i, j) \in G \mid \pi(i) \in C \wedge \pi(j) \in C\}|]$$

Where the expectation is taken over permutations $\pi$ on the vertices of $G$.

As an estimate of $n_1$, we take Euler's formula, which defines an upper bound on the edges in a planar graph:

$$|E(G)| \leq 3\,|V(G)| - 6$$

Since the Delaunay triangulation is planar and a clade cannot have more internal edges than a maximal planar graph of that size, this directly translates to an upper bound on $n_{\text{bb}}$:

$$n_{\text{bb}}(G, C) \leq 3\,|C| - 6$$

We then normalize the join counts according to a min-max scaling for these two values:

$$\alpha_C = \frac{n_{\text{bb}}(G, C) - n_0}{n_1 - n_0}$$

Not that $n_0$ is not really a minimum: in cases of systematically intertwined clades $n_{\text{bb}}$ may even be lower than in the purely random case. This would result in $\alpha_C < 0$. However, we neither expected nor observed negative values for $\alpha_C$ in our experiments.

As a last modification to the score, we want to bring the measure in line with our chosen terminology of clade overlap. $\alpha_C$ measures the compactness of a clade, which is large if the overlap is low. Thus, we invert the score to obtain the overlap score used in our experiments:

$$\text{overlap} = 1 - \alpha_C$$

#### S5.2 Diversity-space dependence

In the ExpSim scenario we model diversification as a result of spatial expansion: speakers in larger areas are less in contact, which leads to language divergence and eventually a split into two distinct languages. This implies that a clade of languages that is expanding faster will also diversify more (i.e. result in more languages). This effect is observable in a correlation between the rate of spatial expansion of a clade and the rate of diversification. Again, the MigSim scenario does not show such a pattern: In random walks (directed or not) the diversification and migration rates may vary, but there is no reason to expect these two to be correlated. We estimate the migration rate in a clade by the spatial variance divided by its age. Similarly, we define the log-diversification rate as the logarithm of the number of leaves divided by the clade age[1]. We measure the correlation between the migration and the log-diversification rate using the the Pearson correlation coefficient and denote this score by *diversity-space dependence*. The diversity-space dependence is expected to be close to 0 if movement and diversification are independent, and tends towards 1 if they are strongly correlated.

#### S5.3 Tree imbalance

Furthermore, we measure the imbalance of the phylogenetic tree. This captures a way in which the previous two observations about the ExpSim scenario – disjoint language areas and space-diversity dependence – interact: Due to disjoint language areas some clades will be restricted by other languages (and geographic constraints) in their spatial spread, while other clades at the frontier of the expansion can continue to grow into new areas. The space dependent diversification in turn means that these locked-in clades will not diversify as much and contain fewer languages. This effect manifests itself in an imbalanced tree. Tree imbalance has been studied extensively in the literature on phylogenetic tree shape [5] and researchers have proposed different scores to measure imbalance. We use the weighted imbalance score $I$ introduced in [6]. This score has an expectation of 0.5 for an unbiased birth-death process and is lower for unexpectedly balanced and higher for unexpectedly imbalanced trees.

#### S5.4 Migrations, expansions, and Bantu

In Figure 5 of the main paper we show all three statistics for the MigSim scenario, the ExpSim scenario, and for the phylogeographic reconstruction of the Bantu expansion. We can see that the

[1]We take the log-diversification rate because the number of leaves is growing exponentially with time, in contrast to the spatial variance, which grows linearly with time (in a random walk scenario).

MigSim scenario produces migrations with high clade overlap, low diversity-space dependence and low tree imbalance compared to the ExpSim scenario. In fact, the diversity-dependence is (widely) distributed around 0 and the tree imbalance falls around 0.5, as expected, since the diversification dynamics and the movement dynamics are independent. In both cases the ExpSim scenario would clearly deviate from such a null hypothesis.

Further, we can see that the Bantu expansion strikingly follows the distribution of the ExpSim scenario. In the Bantu case we computed the statistics on every sample from the posterior distribution, yielding the plotted distribution of the statistics. Since the statistics from the Bantu expansion represent one historical scenario (rather than many simulated ones, as in MigSim and ExpSim), it is not surprising that the distributions are slightly narrower. The important observation is the clearly higher agreement with the ExpSim scenario, suggesting that the Bantu case study aligns with an expansion scenario.

We want to emphasize that we see these statistics as a first step towards a more detailed characterization of historical migration processes and that any such statistic can only be part of a broader discussion of plausibility of reconstructions in a historical context.

#### S6 Effect of missing first clades

We also explored the effect of missing first clades, as it might occur due to language shifts, population shifts or uneven sampling. In the top panel of Figure S6 we render the reconstruction of the Bantu languages, based on the full sample from [7] (including an outgroup of Grassfields languages). The root – marked with a circle – is placed by the reconstruction in central/western Cameroon, close to the hypothesized homeland of Bantu languages. In the bottom panel we show the reconstruction based on a reduced sample, where we removed the outgroup and the first three clades. In line with our arguments about the importance of the first clades (Section 5 (b)), we see that not including these clades has a severe effect on the reconstruction of the homeland. The result is also in line with the convexity property introduced in Section 5 (a) of the main paper.

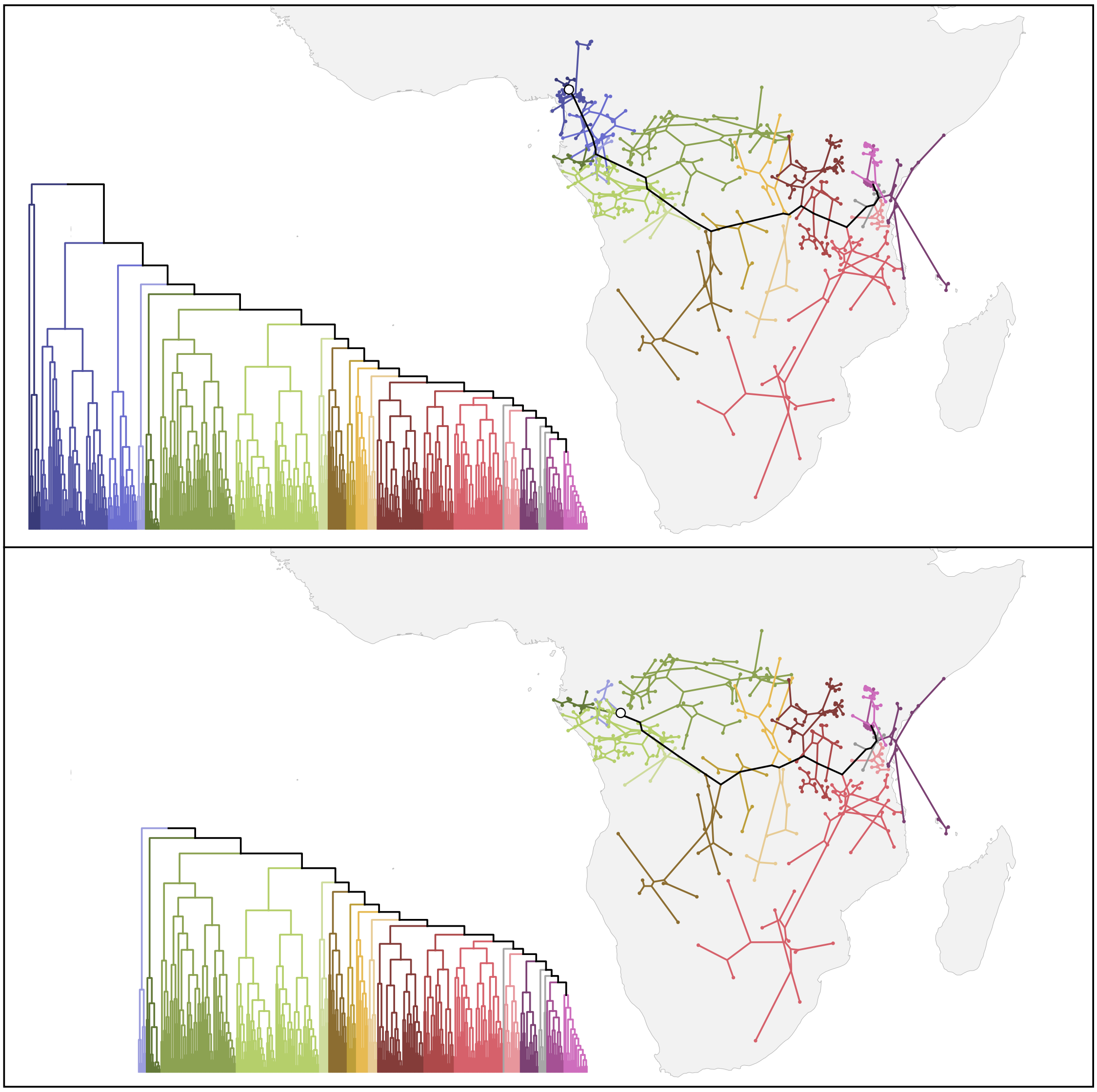

Figure S6: The effect of missing first clades.
